# scTimeBench: A streamlined benchmarking platform for single-cell time-series analysis

**DOI:** 10.64898/2026.03.16.712069

**Authors:** Adrien Osakwe, Eric H. Huang, Yue Li

**Affiliations:** Quantitative Life Sciences Program, McGill University, 550 Rue Sherbrooke O., Montreal, Quebec, Canada, H3A 1B9; School of Computer Science, McGill University, 3480 Rue University, Montreal, Quebec, Canada, H3A 2A7

**Author notes:** These authors contributed equally to this project and are both co-first authors (Adrien Osakwe, Eric H. Huang).

**Keywords:** single cell, time-series, cell lineage, benchmark, scRNA-seq

## Abstract

Temporal modelling of single-cell gene expression is essential for capturing dynamic cellular processes, yet a systematic framework for evaluating time-aware trajectory inference methods has not yet been established. Here, we present a modular and scalable benchmark designed to assess methods across three critical tasks: forecast accuracy (temporal cell alignment) for projecting cells to unseen time points, embedding coherence between original and projected data, and cell-type lineage fidelity. We evaluated nine state-of-the-art methods, which are broadly categorized into 7 forecasting-based and 2 optimal transport (OT)-based methods across eight diverse datasets spanning four species. Our results show that while several methods achieve high forecast accuracy, they often fail to preserve biological signals, both in their latent spaces and in cell lineage reconstruction. Notably, most methods confer low lineage fidelity and often underperform compared to a correlation baseline. We further demonstrate that integrating pseudotime can effectively denoise trajectories by aligning the data snapshots with the intrinsic biological clock in each cell. Finally, to streamline benchmarking for temporal single-cell analysis, we built one of the first self-contained Python packages for the research community: https://github.com/li-lab-mcgill/scTimeBench.

## Introduction

Cells in living organisms arise through a sequence of differentiation processes that transform them from immature, pluripotent states into specialized, mature cell types. To investigate these processes, biologists have generated temporal single-cell RNA-sequencing (scRNA-seq) datasets, which allow for the study of gene expression dynamics (Ding et al., 2022). As current sequencing techniques destroy cells, each cell is limited to a single time point, requiring computational methods for temporal cell alignment. Existing methods leverage either the observed time points or proxies for the internal clock of cells (i.e. pseudotime) to recover developmental trajectories, and reveal insights on tissue development and disease progression (Trapnell et al., 2014; Saelens et al., 2019).

Time-supervised methods are crucial to single-cell trajectory inference (Zhang et al., 2025). Forecasting-based methods leveraging Ordinary Differential Equations (ODEs) and diffusion seek to reconstruct biological processes as continuous trajectories (Zhang et al., 2024; He et al., 2026; Zhang et al., 2025). Optimal Transport (OT)-based methods such as WOT and Moscot have gained prominence for their ability to provide discrete, cell-to-cell mappings across distinct time points (Schiebinger et al., 2019; Klein et al., 2025). Importantly, these supervised temporal models can project cells through time, facilitating new biological discoveries. In particular, these models promise in-silico cellular perturbations via the concept of virtual cells (Roohani et al., 2025), which goes beyond hypothesizing candidate drug targets by elucidating the underlying transcriptional mechanisms susceptible to various drug effects. The advancement of translational medicine relies on robust time-supervised methods whose progress is hindered by the lack of a systematic evaluation framework. While time-independent models have been benchmarked for characterizing cell trajectories, time-dependent assessments over a diverse set of methods remain limited (Inecik et al., 2025). Comparative analyses of time-supervised methods are limited to small-scale custom benchmarking experiments (Zhang et al., 2024, 2025; He et al., 2026). Currently, most existing studies focus on gene expression forecasting, without a comprehensive evaluation of the underlying biology (Zhang et al., 2024; Jiang and Wan, 2024; Jiang et al., 2026). Ultimately, the rapid generation of new datasets and methods calls for a modular, scalable, and adaptable benchmarking framework.

In this study, we introduce `scTimeBench,` a modular and scalable benchmarking framework for cell lineage reconstruction and projection in single-cell time-series data (Figure 1). Our framework provides a practical way for researchers to assess the performance of their methods at three core time-series tasks over a range of datasets and scenarios: (1) the accuracy of gene expression projection over time, (2) the coherence of projected latent spaces and (3) the fidelity of the observed cell lineage of projected cells. Through the evaluation of these tasks, we identify the limitations of forecasting-based and OT-based methods, and their unsolved challenges.

**Figure 1.**
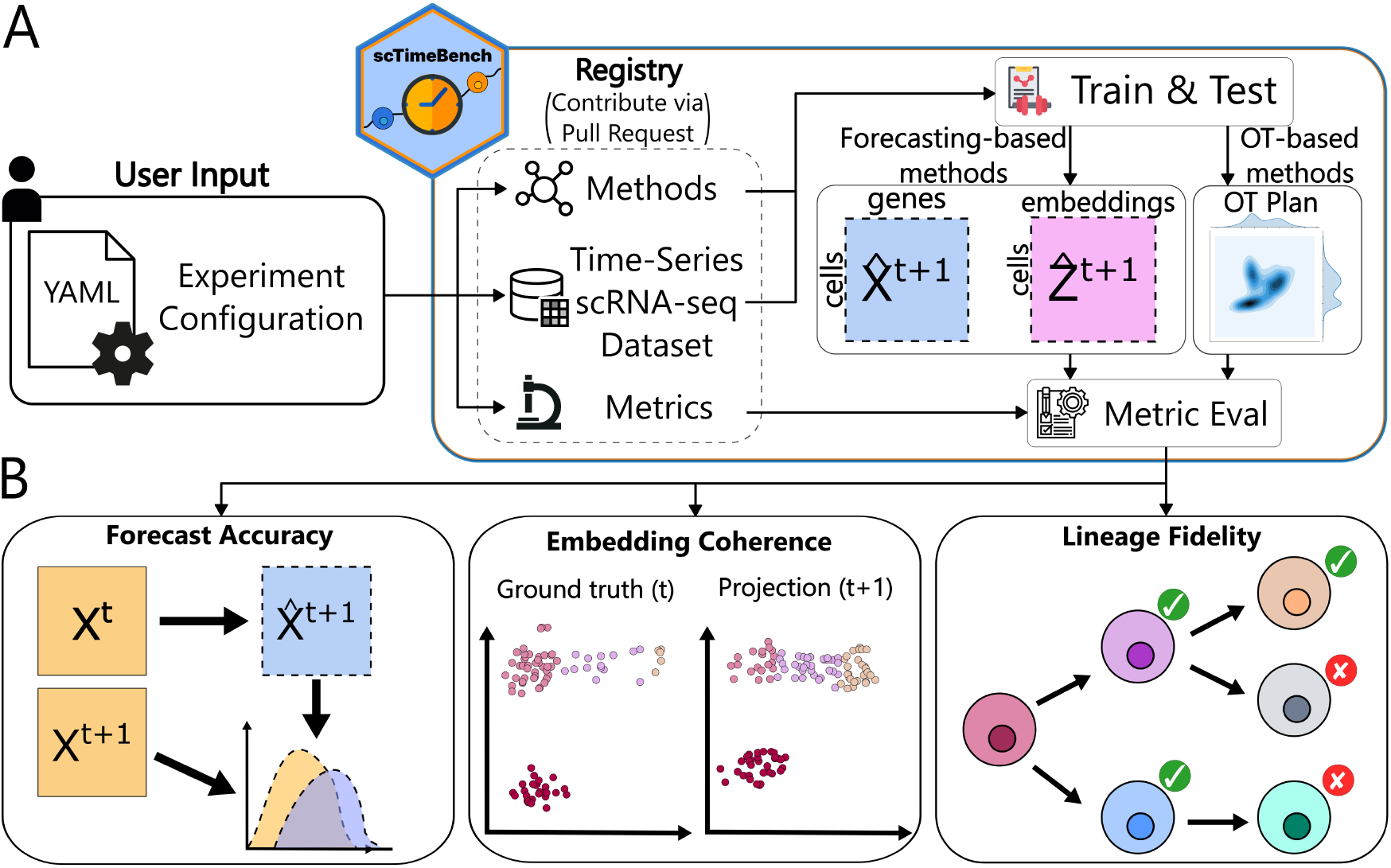
`scTimeBench` **overview** (A) `scTimeBench` provides users with an intuitive environment for method evaluation. Users provide a YAML config file to select a model, datasets, and metrics. (B) Each method was evaluated on forecasting gene expression (forecast accuracy), distilling a meaningful latent space to known cell types (embedding coherence), and recapitulating known cell differentiation pathways (lineage fidelity).

## Materials & Methods

### scTimeBench overview

Our central goal is to systemically evaluate a diverse set of temporal-aware methods that project cells over time in terms of preserving distributional accuracy and biologically meaningful lineage structure. To this end, we designed `scTimeBench` to be modular and scalable to accommodate the growing number of available methods, datasets, and experimental scenarios with minimal effort from users (Figure 1). The pipeline allows users to tailor their own benchmark evaluation setups, configured by custom YAML files, which detail the exact models, datasets, and metrics to use. We apply the framework to eight datasets (Table S1), spanning diverse organisms, tissues, and dataset sizes to benchmark methods across multiple biological and computational scales. The flexibility of our framework ensures that the benchmark can be continuously updated as the field evolves to include newly curated datasets and evaluation scenarios. In the following sections, we describe the three complementary evaluation metrics.

### Forecast accuracy

Forecast accuracy evaluates the alignment of the projected gene expression from an earlier timepoint (*t*) with the observed expression at the subsequent timepoint (*t* + 1). We benchmarked performance on 6 datasets (Table S1) spanning zebrafish (ZB), drosophila (DR), mouse embryonic fibroblasts (MEF) and human developing pancreas (early development with the Ma dataset and late with Olaniru). The ZB, DR, and MEF datasets used the top 2000 highly variable genes whereas Ma and Olaniru were whole transcriptome. A sixth dataset composes of both the Ma and Olaniru datasets to assess robustness to batch effects. For each dataset, we created three scenarios of varying difficulty: an interpolation task (easy), an extrapolation task (medium), and a joint interpolation and extrapolation task (hard). In all scenarios, performance was evaluated on the held-out time points. To quantify forecast accuracy, we used four metrics implemented in the *geomloss* package: Wasserstein Distance (WD), Gaussian maximum mean discrepancy (MMD), Energy distance MMD, and the Hausdorff loss (Feydy et al., 2018). These metrics emphasize complementary evaluations of forecast accuracy: WD quantifies the average cost of moving the projected cells to the ground truth; MMD uses kernel-derived feature mappings (Gaussian and Energy distance) to compare point clouds; Hausdorff highlights the worst-case mismatches in the projections. As the ranges of metrics varies substantially based on method performance, we aggregated results by ranking methods within each scenario, reporting the mean rank across metrics as the final forecast accuracy score.

### Embedding coherence

Complementary to the forecast accuracy, we evaluate to what degree the projected cells preserve cell-type-specific signals from the original data (Figure 3A). We assess this coherence by two metrics. As the first metric, we compare the inferred embeddings for the projected cells 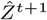 and the actual cells *Z*^*t*^. As the projected cells have no labels, we first use K-nearest neighbors (KNN) to predict their cell types based on the cell-type annotations from the original data. Then, we compute clusters with the Leiden algorithm and use Adjusted Rand Index (ARI) to measure cluster quality. As the second metric, we use a trained classifier to predict cell-type labels of the projected cells. Specifically, we train a random forest classifier on the original data and calculate the average normalized entropy per cell. If the embeddings preserve cell-type-specific signals, we expect low entropy in both the original and projected cells.

### Lineage fidelity

We use three curated datasets from human samples with well-defined cell lineages: (i) the Garcia-Alonso gonad developmental dataset, (ii) the Suo B-cell maturation dataset, and (iii) the Ma early embryonic pancreas developmental dataset (Table S1). After training each model, we project all cells onto the next time point and infer their new cell-type label. For OT-based methods, we use the inferred transport plan to match each cell at time *t* to a cell at time *t* + 1, allowing us to assign a new cell-type label based on the matched target cell. For other methods, we instead annotate the projected cells using CellTypist on the predicted gene expression profiles (Table S2) (Domínguez Conde et al., 2022). We compile all predicted transitions into a cell-type transition matrix and convert it into a binarized lineage graph by applying the threshold that maximizes the total precision and recall (Figure 4A).

To evaluate the accuracy of the predicted lineages, we employ a suite of metrics that quantify both the local (direct cell-type transitions) and global (indirect downstream cell-type) lineage fidelity of the inferred graph. Across all metrics, the reference cell lineage is treated as the ground truth, where known transitions *i* → *j* represent the positive edge. We first assess the cell-type transitions by AUROC and AUPRC to quantify how well each method prioritizes true lineage. In addition, we calculate the Jaccard Similarity between the predicted edges of the binarized lineage graph (*P*) and ground truth edges 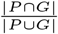.

Additionally, we evaluate lineage fidelity at two resolutions: **single-step** measures the accuracy of immediate developmental transitions; **multi-step** captures broader, long-range connectivity (Figure 4A). Specifically, single-step assesses the exact match between the predicted graph and the ground-truth graph, which can be overly stringent. As a more lenient metric, multi-step computes Floyd-Warshall reachability between two cell types. This rewards the preservation of the overall direction and flow of the cell lineages, even if intermediate cell types are missing. As a baseline method, we infer cell lineages by assigning cell-type transitions with the largest Spearman rank correlation between cells at time t and t+1 in the gene expression space.

### Method implementations

The current version of our benchmark includes 9 methods (Table 1). All included methods were implemented based on the tutorials and documentation provided by the original studies. For methods generating errors at runtime, changes were made to the codebase and documented in a git submodule for reproducibility. As the PHATE algorithm used within MIOFlow does not allow for gene expression reconstruction, we substituted it for PCA as done in previous studies (Zhang et al., 2024). To include PISDE in the embedding coherence analysis, we fit a PCA to the original data and applied it to the projected expression profiles. As the CellMNN codebase was unavailable, we re-implemented its extensions to the MNN package based on the original publication with assistance from a GitHub Copilot agent (Pervez et al., 2024; Bassewitz et al., 2025). The evaluated OT methods cannot predict new cells at unseen time points and are thus excluded from the forecast accuracy and embedding benchmarks.

**Table 1.** Technical specifications and practical implementation metrics of single-cell trajectory inference methods.

| Method | Algorithm | Generative | Latent Space<sup>1</sup> | Runtime (s)<sup>2</sup> | Code changed | Source |
| --- | --- | --- | --- | --- | --- | --- |
| CellMNN | Mechanistic NN (Local ODEs) | Y | Y | 1449 | Y | Bassewitz et al. (2025) |
| MIOFlow | Geodesic VAE + Neural ODE | Y | Y | 1216 | Y | Huguet et al. (2022) |
| Moscot | Optimal Transport | N | N | 436 | N | Klein et al. (2025) |
| PISDE | Neural SDE + Physics-Informed | Y | N | 13270* | Y | Jiang and Wan (2024) |
| PRESCIENT | PCA + Neural SDE | Y | Y | 7538 | N | Yeo et al. (2021) |
| scIMF | Transformer + Neural SDE | Y | Y | 1035 | Y | Jiang et al. (2026) |
| scNODE | VAE + Neural ODE | Y | Y | 2524* | N | Zhang et al. (2024) |
| SquidDiff | Conditional Diffusion Model | Y | Y | 12986 | N | He et al. (2026) |
| WOT | Optimal Transport | N | N | - | N | Schiebinger et al. (2019) |

Abbreviations: NN (Neural Network), VAE (Variational Autoencoder).
<sup>1</sup>Refers to whether the model has a learned latent space, not just inferred from methods such as PCA.
<sup>2</sup>Runtime was measured on the Suo dataset from training and model generation. Methods marked with \* indicate CPU usage due to lack of GPU support or out of memory errors. All GPU benchmarks used RTX 2080Tis. Methods with a - failed to generate outputs.

## Results

### Forecast accuracy

We first evaluated the forecast accuracy using the ZB, DR and MEF datasets based on the top 2000 HVGs per dataset (Figure 2A) (Farrell et al., 2018; Calderon et al., 2022; Schiebinger et al., 2019). Across metrics, scIMF yielded the highest overall performance, with MIOFlow, scNODE, and PI-SDE trailing with similar performance between them. We note that the top three methods all use variations of jointly training a VAE and neural differential equation. scIMF also yielded the highest overall WD and Hausdorff with the second-best performance in Energy distance MMD but poor performance for Gaussian MMD (Figure S1). Squidiff had the highest performance for Gaussian MMD with an overall average rank of 1.0, but performed the worst in all other categories. This underscores the importance of using multiple metrics.

**Figure 2.**
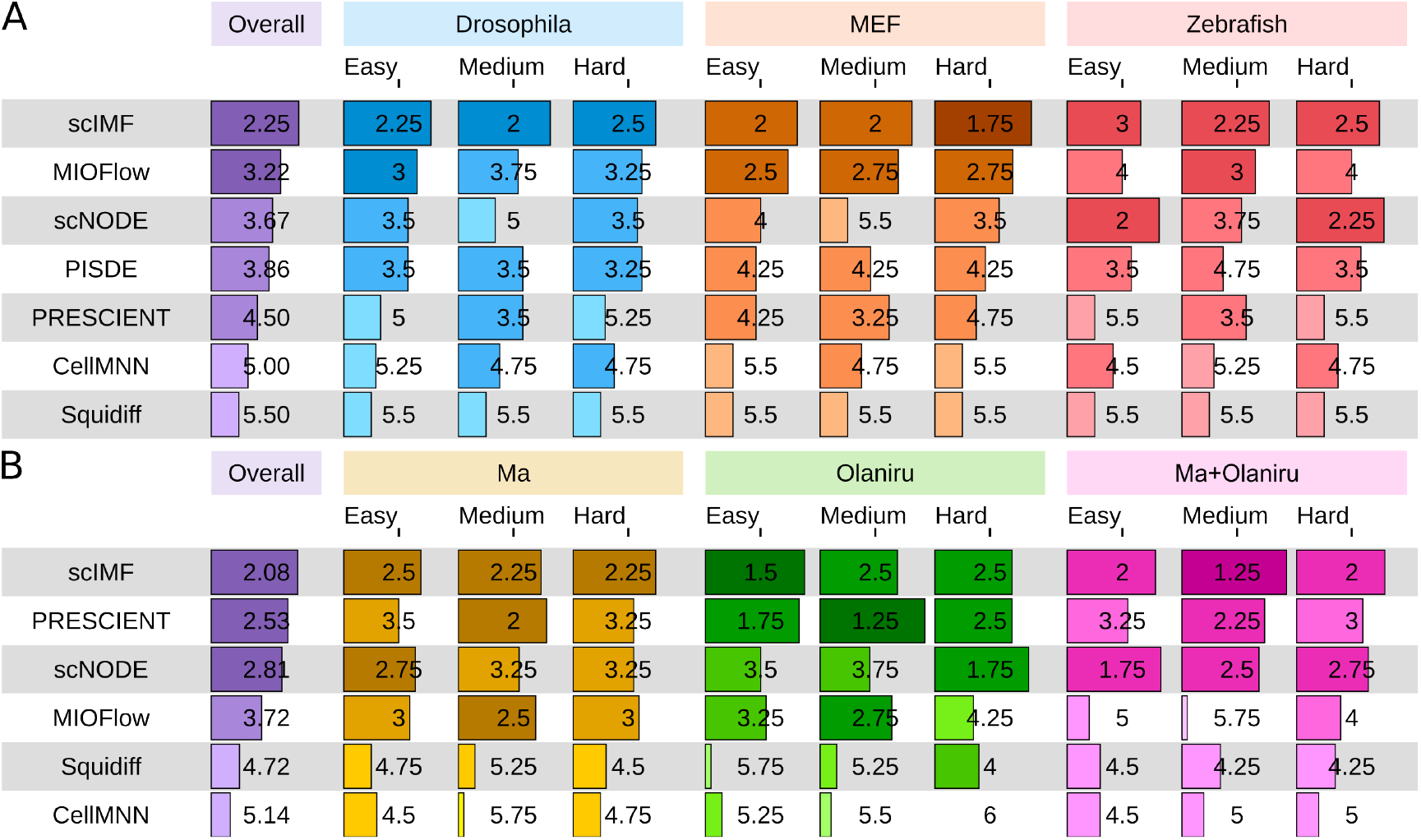
Forecast accuracy. (A) Averaged rank performance on Drosophila, MEF, and Zebrafish datasets. Each method was trained and tested using the HVGs for each dataset. (B) Averaged rank performance on Ma, Olaniru, and combined Ma+Olaniru pancreas datasets using whole transcriptome. PI-SDE was omitted due to out of memory errors.

**Figure 3.**
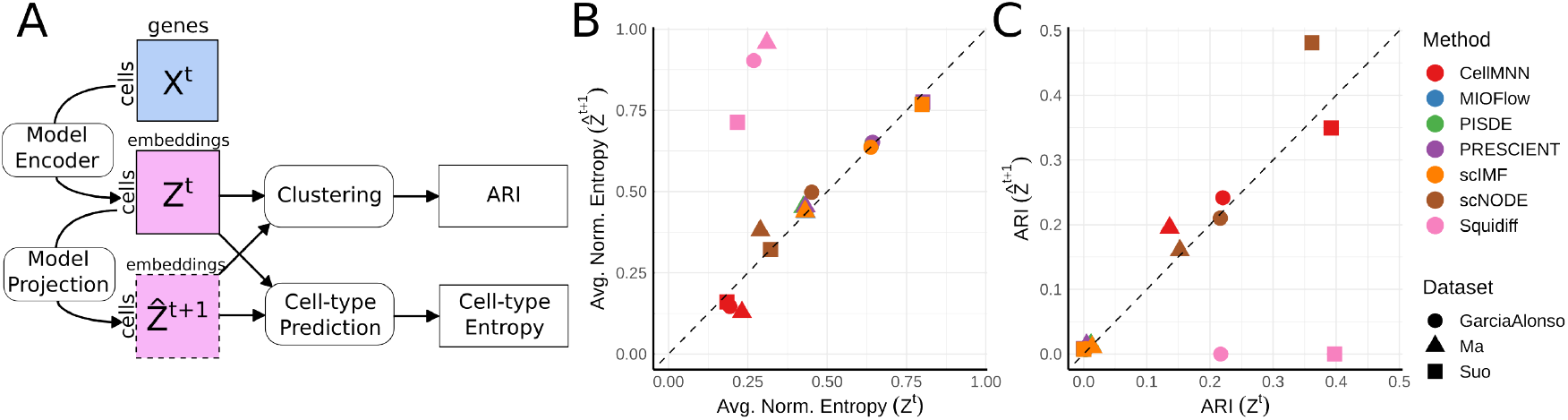
Embedding coherence. (A) Embeddings for ground truth (*t*) and projected cells (*t+1*) are passed through a standard clustering and cell-type prediction pipeline to compute ARI and average classifier entropy. (B) Biological coherence of cell embeddings in terms of classifier entropy (the lower the better). (C) Clustering ARI using embedding *Z*^*t*^ of observed cells at time t and projected embedding for time 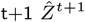.

**Figure 4.**
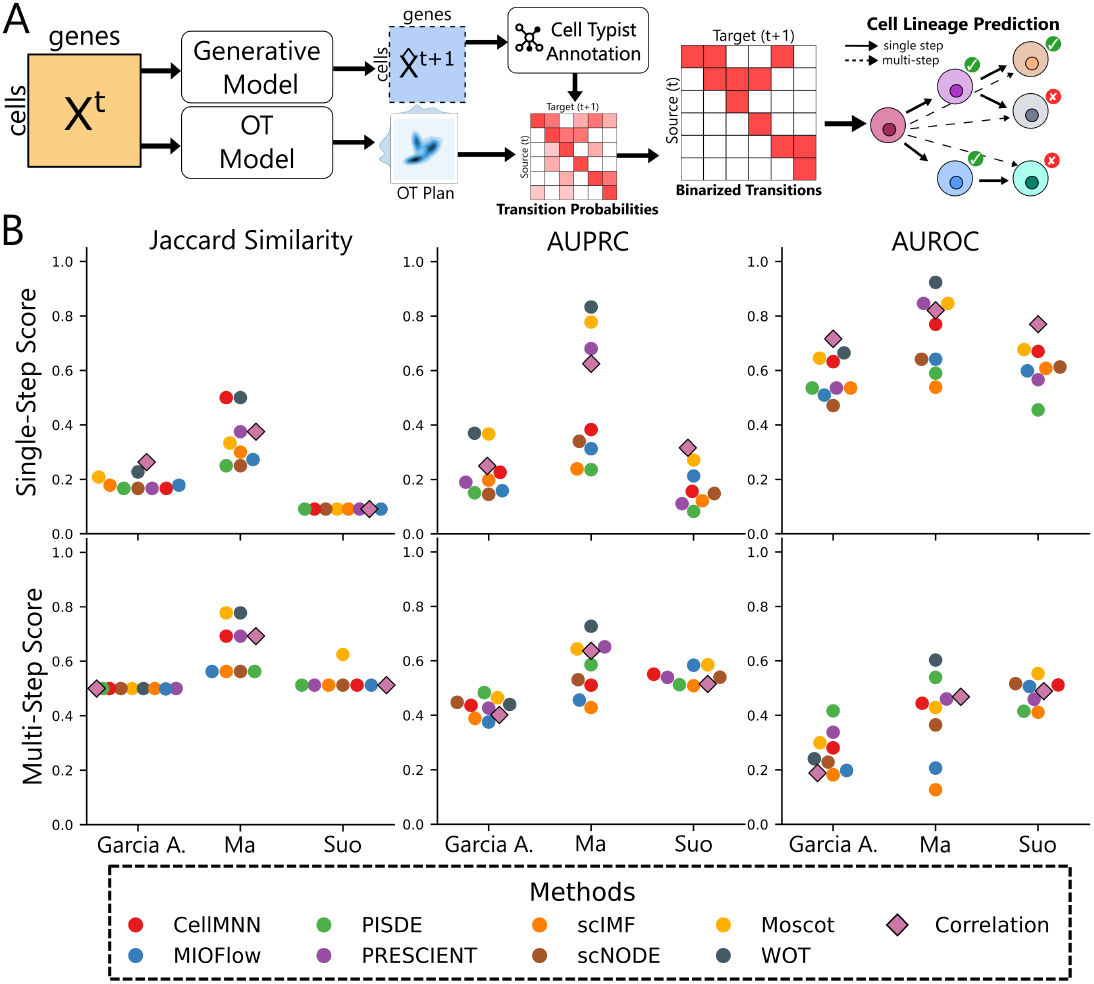
Lineage fidelity benchmarks. (A) Cell lineage prediction pipeline. Cell lineage metrics across datasets for all methods. Squiddiff was omitted due to errors during cell-type inference.

We then evaluated the models on the Ma, Olaniru, and Ma+Olaniru datasets using all genes (Figure 2B) (Ma et al., 2023; Olaniru et al., 2023). As Ma and Olaniru only overlap at three time points, the combined Ma+Olaniru data evaluates the robustness of temporal projections to batch effects and missing data. Once again, scIMF leads with strong performance in WD, Energy MMD, and Hausdorff (Figure S2). While certain methods experienced worse performance on the whole transcriptome and cross-batch scenarios (e.g: MIOFlow), most yielded consistent rankings. In the medium extrapolation task, scNODE and scIMF demonstrate improved rankings when trained on Ma+Olaniru rather than Olaniru, indicating a stronger capacity to integrate multiple batches and leverage larger training sets. Taken together, we identify scIMF as the best forecasting method followed by scNODE.

### Embedding coherence

We assessed biological conservation of the projected cell embedding at the subsequent time points using the Garcia-Alonso, Suo, and Ma datasets (Figure 3A). For each method, we obtained the inferred embeddings of the observed cells *Z*^*t*^ and those of the projected cells 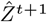 at the same time point. In terms of cell-type entropy (the lower the better), we found that all methods except for Squidiff yielded nearly identical confidence for both groups (Figure 3B). This indicates that the projected embeddings are sufficiently similar to those of the observed cells for the classifier. In contrast, Squidiff yielded much higher entropy for the projected cells, suggesting that their projected embeddings do not align well with their ground-truth embeddings.

In addition, most methods yielded low ARI in both observed and projected cells (Figure 3C). The exceptions were CellMNN and scNODE, which showed higher performance on both groups, while Squidiff showed reasonable ARI performance for observed cells but performed poorly on the projected data. These observations demonstrate that the temporal projections for most methods often yield an embedding space that struggles to distinguish cell types reliably. Together, CellMNN and scNODE conferred the lowest entropy and highest ARI respectively, and therefore preserve the most coherent embedding space.

### Lineage fidelity

#### Cell lineage predictions based on observed time

To evaluate lineage fidelity, we projected cells forward to the subsequent timepoint. These predictions were then aggregated into a cell-type transition matrix to infer a lineage graph (Figure 4A; Section 2). We compared inferred and known lineages using Jaccard similarity, AUPRC, and AUROC (Figure 4B) at two resolutions: single-step (direct transitions) and multi-step (all connected paths). We then computed Jaccard Similarity using method-specific thresholds that maximize the total precision and recall, representing the optimal lineage recovery for each method. Despite this optimization, Jaccard Similarity scores are low especially in the single-step regime. In contrast, several methods achieved higher AUPRC and AUROC. These result suggest that while current methods can effectively rank potential transitions, they struggle to resolve a decisive, sparse lineage structure. Overall, OT-based methods (Moscot, WOT) conferred the best performance across datasets and metrics, with particularly high AUPRC on the Ma dataset (Figure 3). On the other hand, the simple correlation baseline compared competitively against many SOTA methods. This underscores the challenge for obtaining representations that resolve fine-grained directional transitions from single-cell time-series data.

#### Cell lineage predictions based on pseudotime

Our previous results suggest that observed time points can be a noisy proxy for developmental progression, which can obscure the temporal signal needed to learn differentiation dynamics. To probe this, we investigated if using *pseudotime* (Section S1) could improve lineage inference by ordering cells along a continuous developmental axis (Wolf et al., 2018; Haghverdi et al., 2016). Overall, pseudotime improved both multi-step and single-step lineage recovery, with the strongest gains on Garcia-Alonso (Figure 5). In contrast, Ma showed mixed responses, with observed time labels performing better in some settings, suggesting that the benefit of pseudotime depends on how well the developmental ordering of cells is being captured.

**Figure 5.**
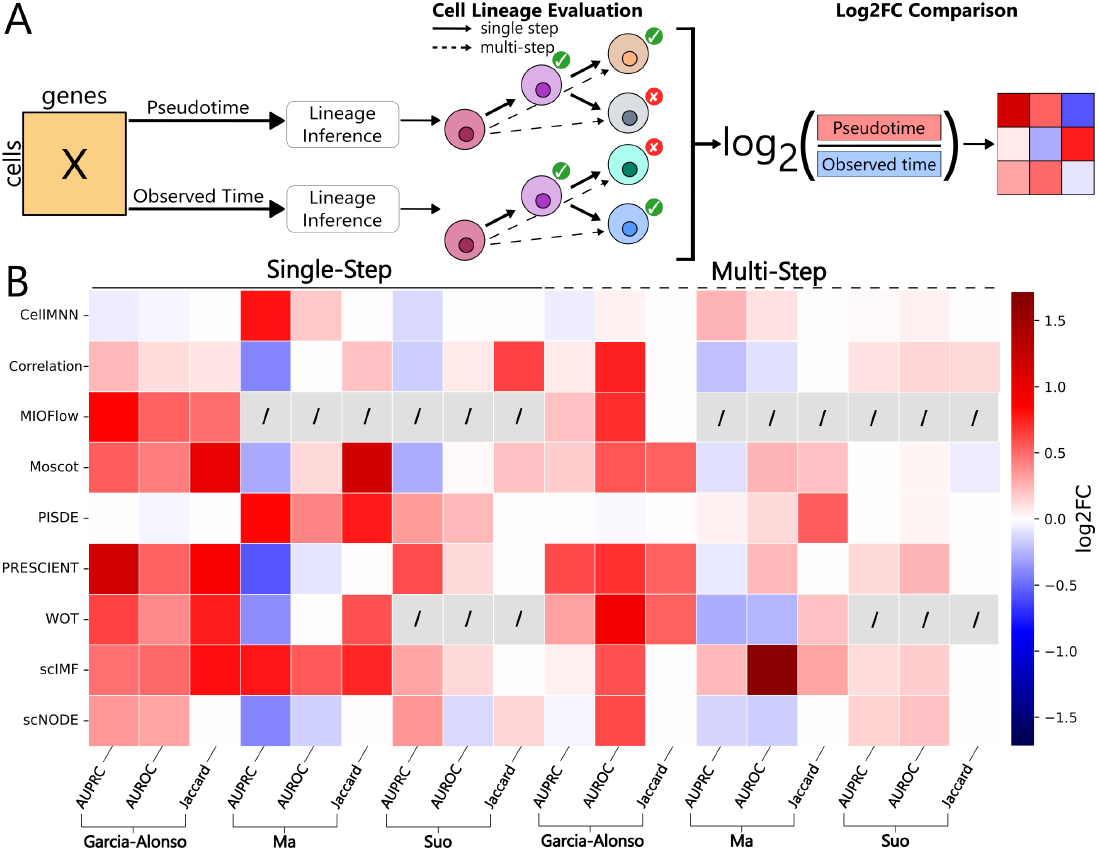
Lineage fidelity: Pseudotime vs. Observed time based inference. (A) We train each method using observed time and pseudotime (Wolf et al., 2018), and evaluated for cell-type lineage recovery in direct (single-step) and downstream (multi-step) link settings. Relative performance across metrics was computed as a log2-fold change. (B) Heatmap of log-fold change between pseudotime and observed time performance. Crossed-out entries indicate failures caused by cell annotation errors or out-of-memory issues.

To diagnose these differences, we visualized the temporal cell-type distribution of observed and projected cells using observed time and pseudotime (Figure 6). We found that the observed cells in the Garcia-Alonso dataset exhibited irregular cell-type mixing across observed time points. For example, primordial germ cell (PGC) proportions drastically decreased at timepoint 12 and then rose up to timepoint 15 before another sharp drop at time point 16. Later states (pre-oocytes/oocytes) are also inconsistently represented across subsequent timepoints. This variability obscures the expected gradual shift from PGCs towards mature oocyte populations. In contrast, the pseudotime-aligned proportions showed a much more coherent progression of cell type proportions, with PGC and germ cell (GC) proportions gradually decreasing to zero and being replaced with more mature cell states. These observations may reflect sampling variability across observed time points, which pseudotime can partially mitigate by aligning cells along a continuous axis. We then examined the cell-type distributions generated by projecting cells with scNODE (Figure 6). The distribution generated from training on observed time points was unable to capture the full cell lineage, stalling at oogonia rather than reaching oocytes; under pseudotime, projections successfully extend to oocytes, though pre-oocytes remain under-recovered. Together, these observations support the hypothesis that temporal label noise and sampling bias in the observed time points can limit inference, and that pseudotime can partially mitigate this by extracting clearer temporal distributions.

**Figure 6.**
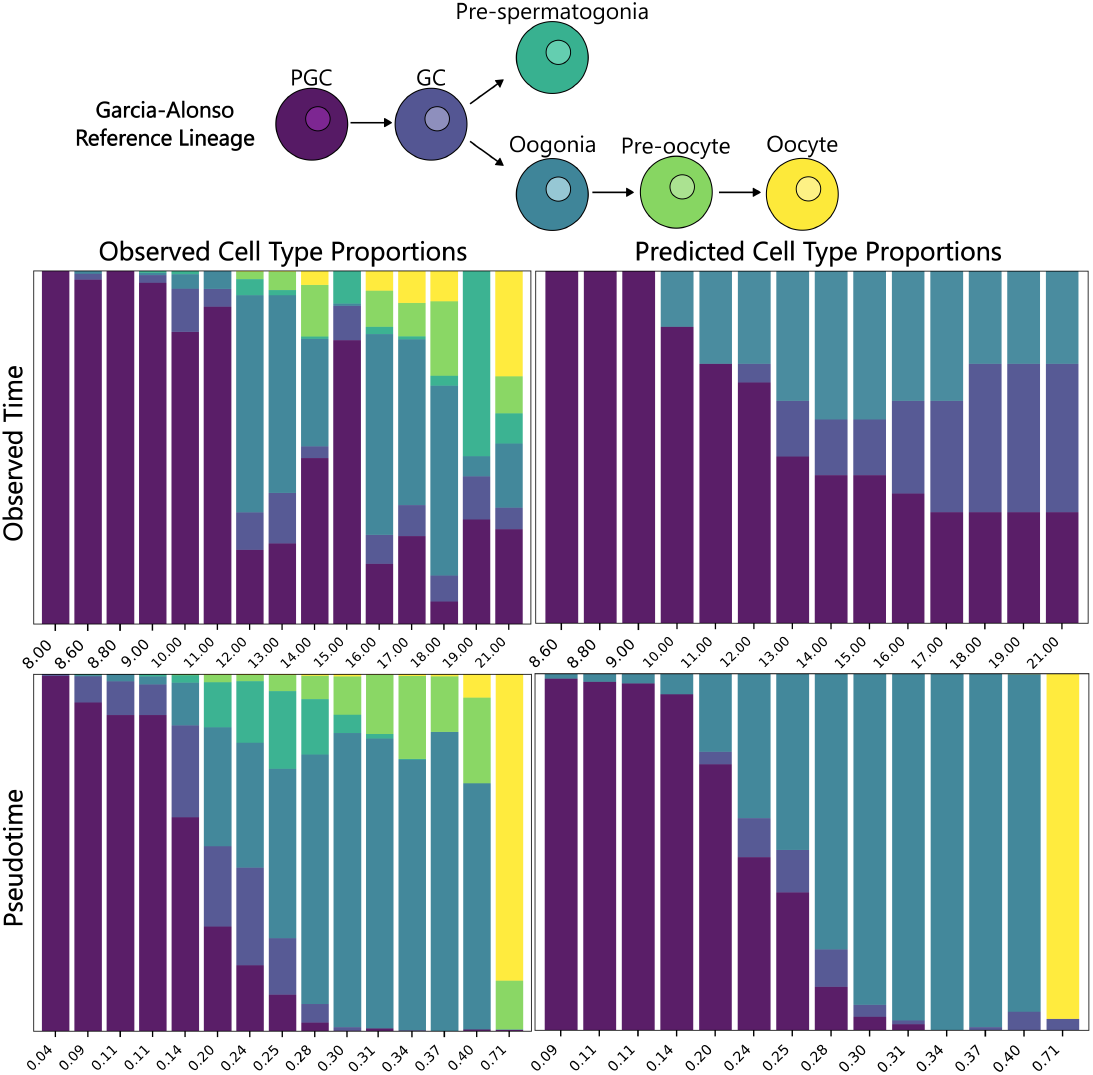
Observed vs Pseudotime time cell-type proportions for observed and predicted Garcia-Alonso data. scNODE cell-type proportions were generated by projecting cells from the initial observed timepoint forward to all subsequent timepoints.

To test whether this pattern generalizes, we repeated the analysis with the Ma dataset, which showed better performance with observed time than pseudotime for scNODE (Figure S3). In this dataset, the temporal distribution of cells in observed time is more consistent with the underlying cell lineage and experiences less variability than Garcia-Alonso, leaving less room for pseudotime to denoise temporal signals. To quantify these shifts, we computed the weighted average cell-type entropy per time bin (Table S3) as a measure of within-bin heterogeneity. Garcia-Alonso shows the largest entropy decrease under pseudotime (more stage-specific bins), whereas Ma shows increased entropy, mirroring their respective improvements and degradations in lineage inference. Thus, some methods can benefit from using pseudotime, which can denoise the temporal distribution of cell types, but the improvement is not guaranteed if the observed time can capture the coherent progression.

#### scTimeBench platform

We implemented `scTimeBench,` one of the first systematic evaluation platforms for temporal single-cell methods. The framework is designed for seamless extensibility, as integrating a new dataset requires only a YAML configuration file to define its attributes; adding a new method involves providing a Python script and supporting bash script; and new metrics are implemented via a standalone Python script (Figure S4). These components are then orchestrated through a single comprehensive YAML configuration file, enabling users to execute `scTimeBench` with just a few lines of code (Figure 7). This will greatly facilitate unbiased method evaluation and new method development in tackling the above challenges.

**Figure 7.**
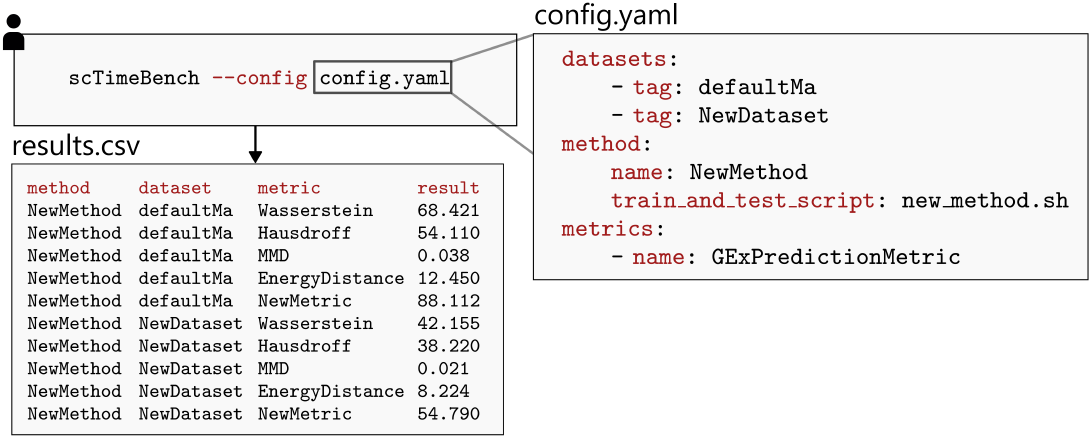
scTimeBench example usage and output.

## Discussion

In this study, we presented a scalable and modular benchmarking framework for the evaluation of temporal scRNA-seq methods. Across eight datasets, we assessed nine methods representing a range of widely used methods. We focused on three core tasks: projecting cells to unseen timepoints (forecast accuracy), preserving biologically meaningful latent space (embedding coherence) and recovering cell differentiation trajectories (lineage fidelity). Together, these analyses highlight both the current strengths of temporal methods and the key challenges that must still be addressed to achieve reliable modelling of cellular dynamics.

Our forecast accuracy results demonstrated that scIMF, through its attention-based SDE architecture, delivered the best overall performance, particularly on whole transcriptome and multi-batch datasets (Figure 2). The joint VAE-ODE framework provided by scNODE also offers a reliable alternative in cases where scIMF fails. Our results also emphasize the importance of evaluating methods across multiple metrics. For example, Squidiff performed the best under Gaussian MMD on the HVG datasets, yet ranked among the worst methods for other forecasting metrics. These findings show that robust benchmarking requires a broader complementary set of evaluations.

Our embedding coherence analyses revealed that CellMNN and scNODE were the only methods able to project cells without substantially disrupting the underlying latent space (Figure 3). In contrast, most methods failed to maintain cluster performance and classifier confidence in both projected and observed cell embeddings. Therefore, despite apparently strong forecast performance, many current methods do not preserve the cell-type-specific signals.

This disconnection was most clearly reflected in our third lineage fidelity analyses. Most methods struggled to recover the correct trajectories when projecting cells forward in time, performing no better than the correlation baseline (Figure 4). While OT-based methods such as WOT and Moscot performed best in lineage reconstruction, their absolute performance was still limited. Therefore, lineage preservation is one of the central weaknesses of the current methods and argue for lineage-based evaluation as a necessary component of future benchmarks.

The improved performance observed under pseudotime-based modelling points to a fundamental limitation of current methods: a mismatch between the temporal distribution of cell types in the data and their true biological progression. Because observed time points are shaped by sampling bias and technical noise, the resulting cell-type distributions do not always reflect the underlying developmental trajectory. As a result, models trained directly on observed time may infer incorrect lineage relationships and generate biologically implausible projections. In contrast, pseudotime provides a way to capture a cell’s internal clock, facilitating biologically meaningful inference. In the Garcia-Alonso dataset, where pseudotime modelling improved performance, the observed-time cell-type distributions were notably discordant with the inferred developmental ordering (Figure 6). In contrast, the Ma dataset showed more biologically consistent cell-type distributions across observed time points, and accordingly did not exhibit the same consistent gains under pseudotime. Together, our findings suggest that future progress will depend on methods that do not treat observed time as the sole supervisory signal, but instead integrate it with trajectory-informed representations such as pseudotime.

Finally, the modular design of our framework positions it as a adaptive benchmark platform for temporal single-cell modelling. As new approaches emerge to better reconcile observed time with pseudotime, `scTimeBench` provides a systematic platform for determining which tools reliably integrate chronological dynamics with biological trajectories to generate meaningful cell projections. By supporting the continued addition of new methods, datasets, and evaluation metrics, `scTimeBench` will evolve alongside the field and remain a relevant resource for method development and comparison.

## Acknowledgements

Y.L. is supported by Canada Research Chair (Tier 2) (CRC-2021-00547), NSERC Discovery Grant (RGPIN-2016-05174), and CIHR Project Grant (PJT-540722). A.O. is supported by the CIHR Canada Graduate Research Scholarship - Doctoral award.

## Data and code availability

All datasets used in this study were acquired from publicly available sources. The processed .h5ad files used in the analysis will be added to Zenodo upon publication. The benchmarking framework presented in this study and all associated analysis scripts are publicly available on GitHub: https://github.com/li-lab-mcgill/scTimeBench/.

## Supplementary Materials

### S1 Pseudotime inference methodology

To enable pseudotime-based analyses, we employed *Diffusion Pseudotime* as implemented in the scanpy framework (Wolf et al., 2018; Haghverdi et al., 2016). This method infers pseudotime as a continuous metric of cellular progression by calculating the relative distances of cell embeddings across a low-dimensional space of a given scRNA-seq dataset. Diffusion pseudotime requires a designated root to anchor the trajectory; we consistently defined this starting point as the first cell captured of the earliest time point. The resulting pseudotime of any given cell represents its distance from this root within the inferred low-dimensional space. We then discretize the data into *n* pseudotime bins (*n* = 15 for the Garcia-Alonso and Suo datasets, and *n* = 10 for the Ma dataset). Cells were assigned to the nearest bin while ensuring an approximately uniform distribution of cells across each interval. The resulting labels provide an approximation of cellular progression relative to the progenitor cell.

### S2 Supplementary Tables

**Table S1.** Summary of benchmark datasets including organism, tissue, and feature counts.

| Dataset | Organism | Tissue | Cells | #tps | Genes | Source |
| --- | --- | --- | --- | --- | --- | --- |
| ZB | Zebrafish | Embryo | 38,731 | 12 | 2,000 | Farrell et al. (2018) |
| DR | Drosophila | Embryo | 27,386 | 11 | 2,000 | Calderon et al. (2022) |
| MEF | Mouse | Embryo | 236,285 | 19 | 2,000 | Schiebinger et al. (2019) |
| Ma (Forecast) | Human | Pancreas | 76,806 | 8 | 29,534 | Ma et al. (2023) |
| Olaniru | Human | Pancreas | 26,220 | 7 | 29,534 | Olaniru et al. (2023) |
| Ma+Olaniru | Human | Pancreas | 103,026 | 15 | 29,534 | Ma et al. (2023); Olaniru et al. (2023) |
| Garcia-Al. | Human | Gonad | 10,993<sup>1</sup> | 15 | 27,950 | Garcia-Alonso et al. (2022) |
| Ma (Lineage) | Human | Pancreas | 6,312<sup>1</sup> | 8 | 29,534 | Ma et al. (2023) |
| Suo | Human | B cells | 65,413<sup>1</sup> | 11 | 33,538 | Suo et al. (2022) |

<sup>1</sup>Cell counts after filtering for cell types which exist in the reference lineage.

**Table S2.**
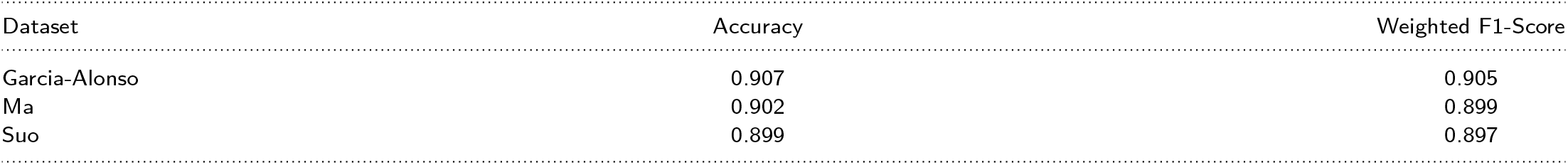
Cell Typist average 3-fold cross-validation scores. We used CellTypist to classify the projected gene expression for generative models.

| Dataset | Accuracy | Weighted F1-Score |
| --- | --- | --- |
| Garcia-Alonso | 0.907 | 0.905 |
| Ma | 0.902 | 0.899 |
| Suo | 0.899 | 0.897 |

**Table S3.** Weighted average entropy of cell types per timepoint.

| Dataset | Real Time Entropy | Pseudotime Entropy |
| --- | --- | --- |
| Garcia-Alonso | 1.203 | 0.760 |
| Ma | 0.651 | 0.702 |
| Suo | 1.967 | 1.646 |

### S3 Supplementary Figures

**Figure S1.**
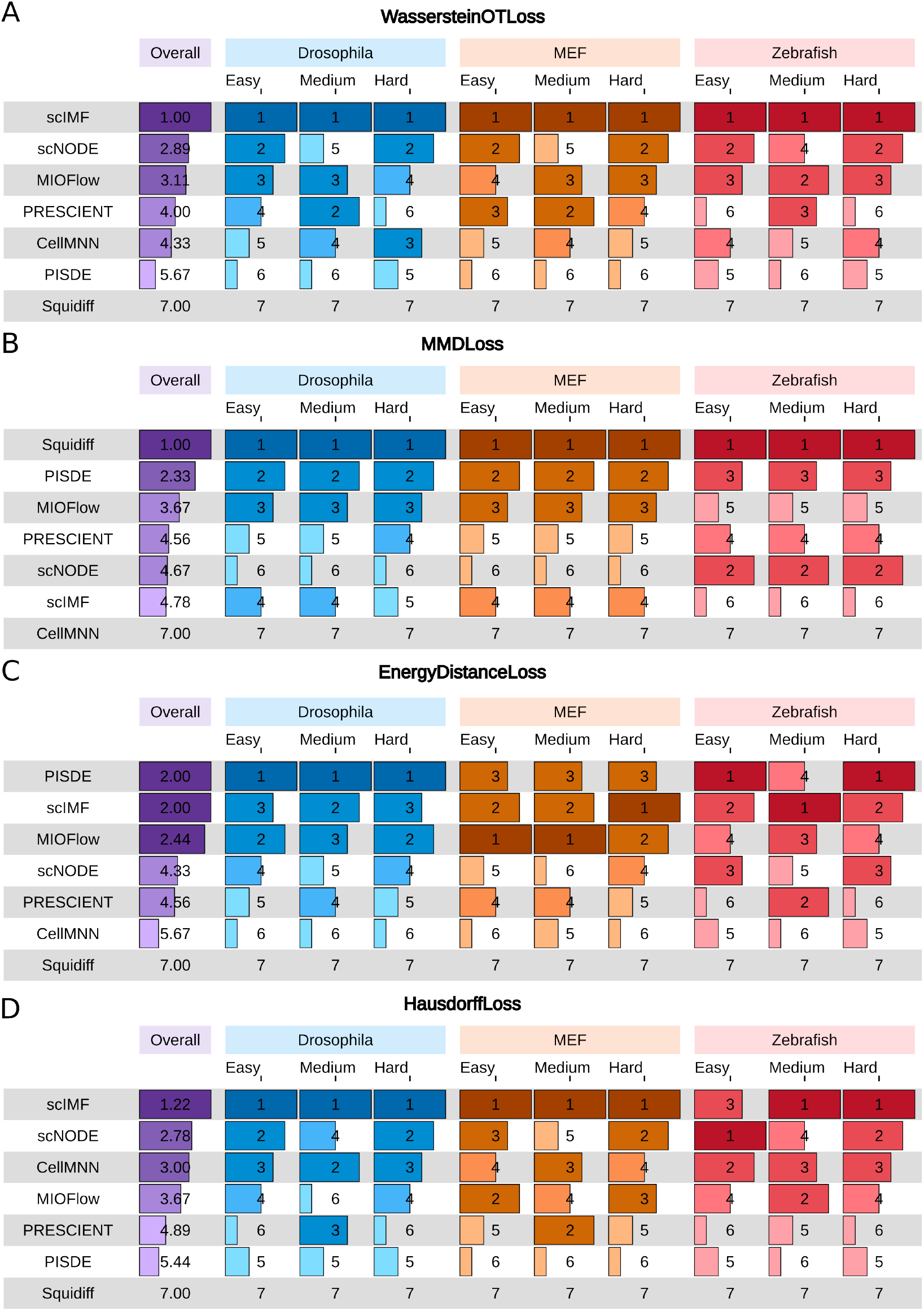
Average rank metric for HVG datasets in terms of Wasserstein distance (A), Gaussian MMD (B), Energy Distance MMD (C) and Hausdorff Loss (D).

**Figure S2.**
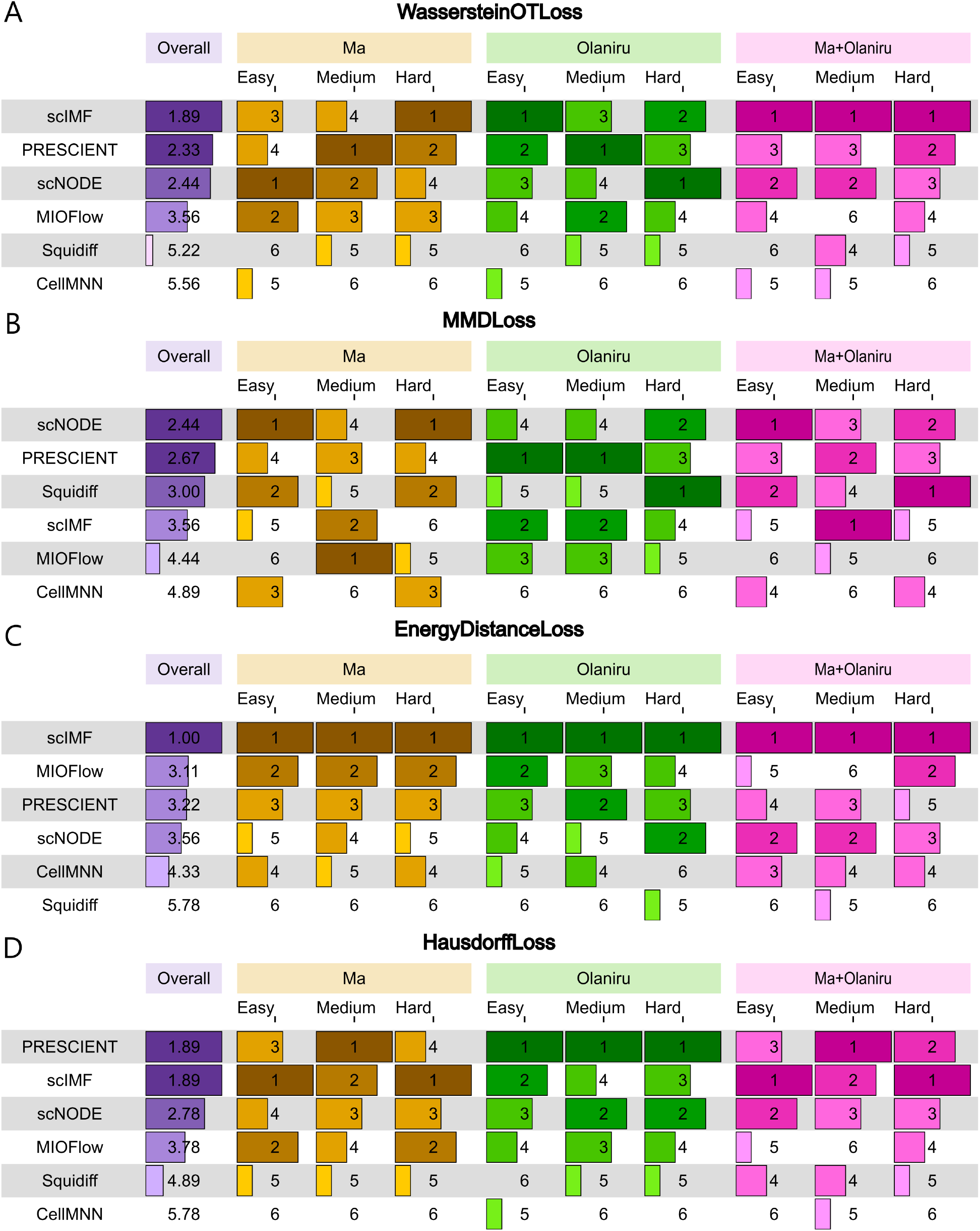
Average rank metric for pancreas development datasets in terms of Wasserstein distance (A), Gaussian MMD (B), Energy Distance MMD (C) and Hausdorff Loss (D).

**Figure S3.**
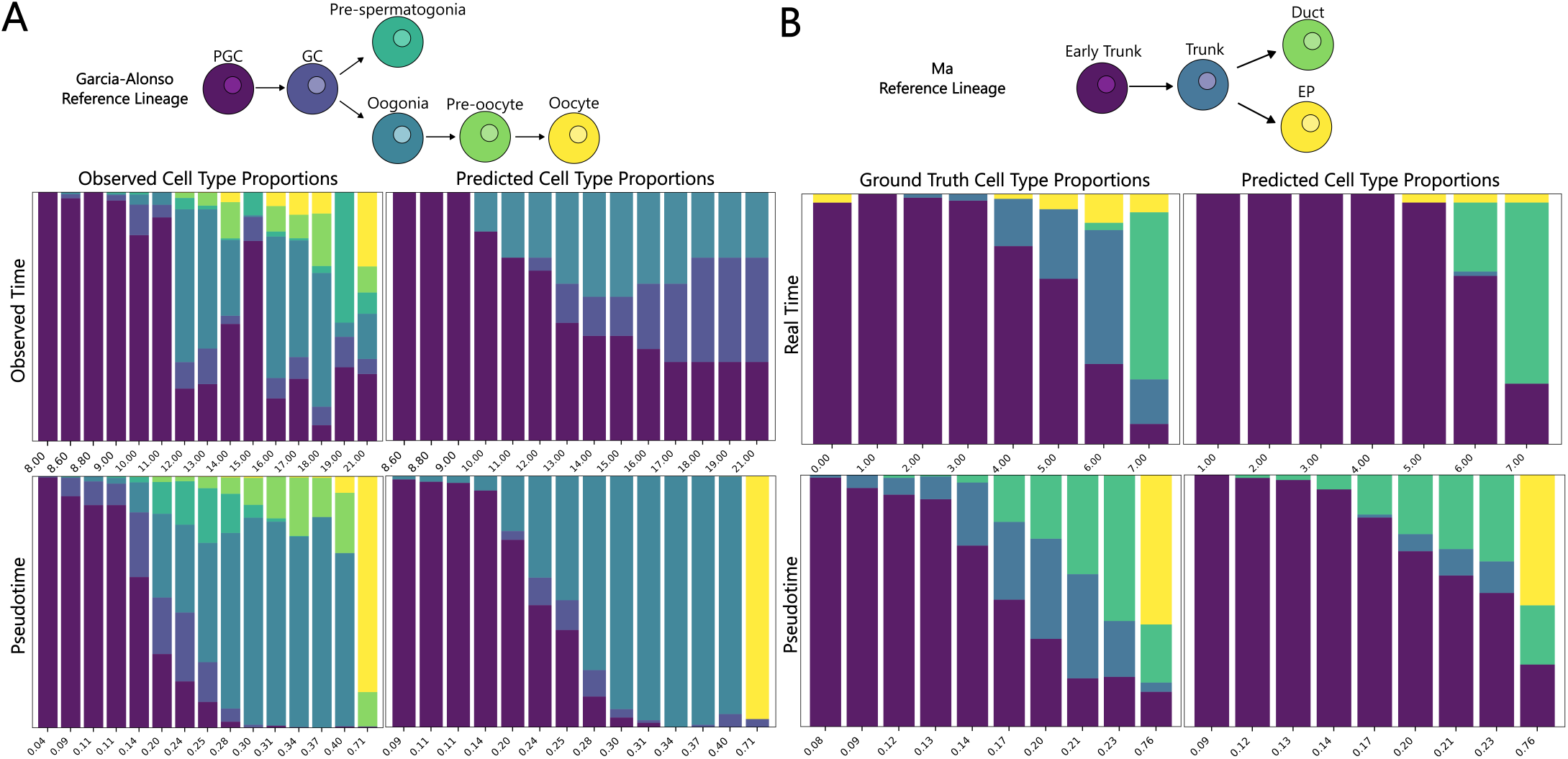
Pseudotime vs. Observed time cell-type proportions for observed and predicted Garcia-Alonso (A) and Ma (B) data. scNODE cell-type proportions were generated by projecting cells from the initial observed timepoint forward to all subsequent timepoints.

**Figure S4.**
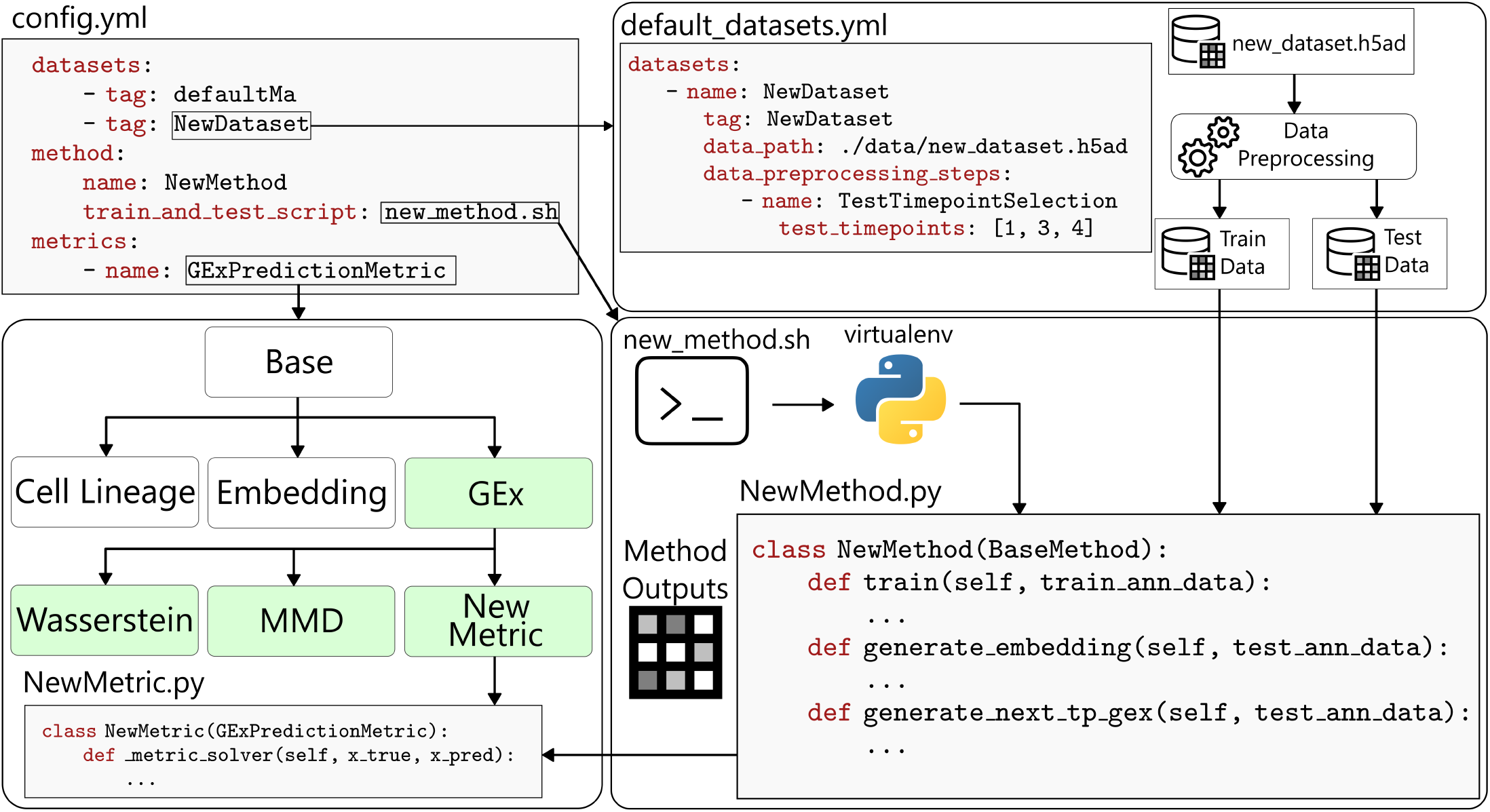
scTimeBench pipeline. The framework utilizes a centralized configuration file to orchestrate the execution of datasets, models, and metrics. **Dataset Integration:** New datasets require a standard Anndata object (.h5ad) file, and defined preprocessing routines for consistent train-test splits. **Model Integration:** The platform supports diverse model architectures, requiring users to provide (1) a shell script that specifies a virtual environment and (2) a Python implementation of the model training and output generation. **Metric Integration:** Metrics are organized into a class-based hierarchy to streamline comparative analysis. For example, all Gene Expression (GEx)/forecast accuracy metrics inherit a unified interface, requiring only the definition of an evaluation function that accepts observed GEx *X*^*t*^ and predicted GEx 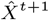. Specifying a parent class (e.g. GExPredictionMetric) in the configuration will run all existing GEx metrics, including any user-defined custom metrics.

